# A single-nucleus transcriptomic atlas of medium spiny neurons in the rat nucleus accumbens

**DOI:** 10.1101/2024.05.26.595949

**Authors:** Benjamin C. Reiner, Samar N. Chehimi, Riley Merkel, Sylvanus Toikumo, Wade H. Berrettini, Henry R. Kranzler, Sandra Sanchez-Roige, Rachel L. Kember, Heath D. Schmidt, Richard C. Crist

## Abstract

Neural processing of rewarding stimuli involves several distinct regions, including the nucleus accumbens (NAc). The majority of NAc neurons are GABAergic projection neurons known as medium spiny neurons (MSNs). MSNs are broadly defined by dopamine receptor expression, but evidence suggests that a wider array of subtypes exist. To study MSN heterogeneity, we analyzed single-nucleus RNA sequencing data from the largest available rat NAc dataset. Analysis of 48,040 NAc MSN nuclei identified major populations belonging to the striosome and matrix compartments. Integration with mouse and human data indicated consistency across species and disease-relevance scoring using genome-wide association study results revealed potentially differential roles for MSN populations in substance use disorders. Additional high-resolution clustering identified 34 transcriptomically distinct subtypes of MSNs definable by a limited number of marker genes. Together, these data demonstrate the diversity of MSNs in the NAc and provide a basis for more targeted genetic manipulation of specific populations.

## Introduction

The nucleus accumbens (NAc) plays important roles in drug-taking and -seeking behaviors ^1-3^. The two primary output pathways of the NAc are GABAergic medium spiny neurons (MSNs) that express either dopamine type 1 receptors (D1R-expressing MSNs) or dopamine type 2 receptors (D2R-expressing MSNs)^4^. In rodents, studies show that addictive drugs, including opioids, cocaine, and methamphetamine, elicit differential responses in D1R-and D2R-expressing MSNs ^5-10^. Furthermore, these two striatal MSN populations may have opposing functional roles in reward-related behaviors: whereas activation of D1R-expressing MSNs *increases* compulsive drug intake, activation of D2R-expressing MSNs *decreases* drug reinforcement ^7,9,11,12^. For example, inactivation of D1R-expressing MSNs or activation of D2R-expressing MSNs attenuates heroin seeking, results that support bidirectional modulation of drug-mediated behaviors by distinct MSN cell populations ^8^.

The NAc is broadly divided into two main subregions: the core and shell. These subregions have unique functional roles in motivated behaviors including drug taking and seeking ^13-15^. MSNs in the NAc core and shell differ both in their afferent ^16^ and efferent ^17,18^ projections, as well as their roles in motivated behaviors ^19,20^. The entire striatum, including the NAc, also contains two distinct compartments: the matrix, forming the bulk of the striatum, and the patch-like striosome interspersed throughout the matrix ^21^. These two compartments are also characterized by different afferent and efferent projections ^21^ and, like the core and shell, play unique roles in addiction-like behaviors ^22,23^. The prevailing evidence suggests that the relevance of MSNs to substance use disorders is not fully captured by the standard classifications of D1R/D2R expression and supports a model wherein subgroups of D1R-and D2R-expressing MSNs have distinct roles in addiction-related phenotypes.

Recent single-cell transcriptomic analyses of the NAc have identified distinct subpopulations of MSNs, including groups representing MSNs from the core, shell, matrix, and striosome. Unique gene expression profiles were discovered in MSNs from the matrix and striosome of two rhesus macaques ^24^ and a study of humans identified 10 MSN subpopulations, with the vast majority of cells falling into the primary D1R-or D2R-expressing MSN clusters ^25^. Studies of the NAc from mice and rats also identified several D1R-expressing and D2R-expressing MSN subtypes, as well as additional MSN populations with other genetic markers ^26-30^. These novel MSN subpopulations may have unique roles in addiction-like behaviors ^27-30^, similar to the functional and gene expression differences between matrix and striosomal MSNs. Similarly, a recent examination of dorsal striatum from humans and non-human primates showed the presence of D1R-and D2R-expressing MSNs, with populations of matrix and striosome neurons, and these populations had sex-and cell type-specific differences in gene expression in the context of substance use ^31^. Thus, a more granular classification of MSN populations is needed to better understand individual MSN cell types and their roles in striatal function and motivated behaviors.

Large single-cell transcriptomic studies can be used to define uncommon and novel neuronal populations in nuclei throughout the brain ^26,32-34^. However, a comprehensive cell type-specific atlas of the NAc has not been generated to date. In this study, we make use of our recently-described, largest available single-cell transcriptomic NAc rat dataset (n=96,627 total nuclei) to present high resolution subclustering of MSNs (*n*=48,040 MSN nuclei) ^30^. We identified 34 transcriptomically-distinct cell populations, including previously unidentified subtypes of D1R-and D2R-expressing MSNs. We replicated these findings by identifying the same novel MSN subtypes in an independent NAc rat dataset (*n*=7,641 MSN nuclei) ^27,28^. Expression patterns for glutamate, GABA, and acetylcholine receptor genes are presented to further phenotype these subclusters. Finally, MSNs from rat, mouse ^26^, and human ^25^ were integrated together and cells scored using genome-wide association study (GWAS) summary statistics for substance use disorder (SUD) phenotypes ^35-37^, revealing potentially differential roles for MSN subpopulations in alcohol use disorder (AUD), alcohol consumption (AUDIT-C), opioid use disorder (OUD), and tobacco use disorder (TUD).

## Results

### Identification of five major MSN cell types in the rat nucleus accumbens

Single nucleus transcriptomic data from male, drug-naïve Brown Norway rats (*n*=11) were aligned to the rat genome and cleaned to remove low-quality nuclei and putative doublets, yielding 96,627 nuclei (Figure 1A). Clustering revealed the presence of a large cell population expressing high levels of *Bcl11b* and *Pde10a*, two known markers of MSNs (Figure 1B-C). Additional marker analyses revealed that the dataset also contained populations of GABAergic interneurons, cholinergic neurons, and all expected major types of glial cells (Figure 1D-K).

**Figure 1:**
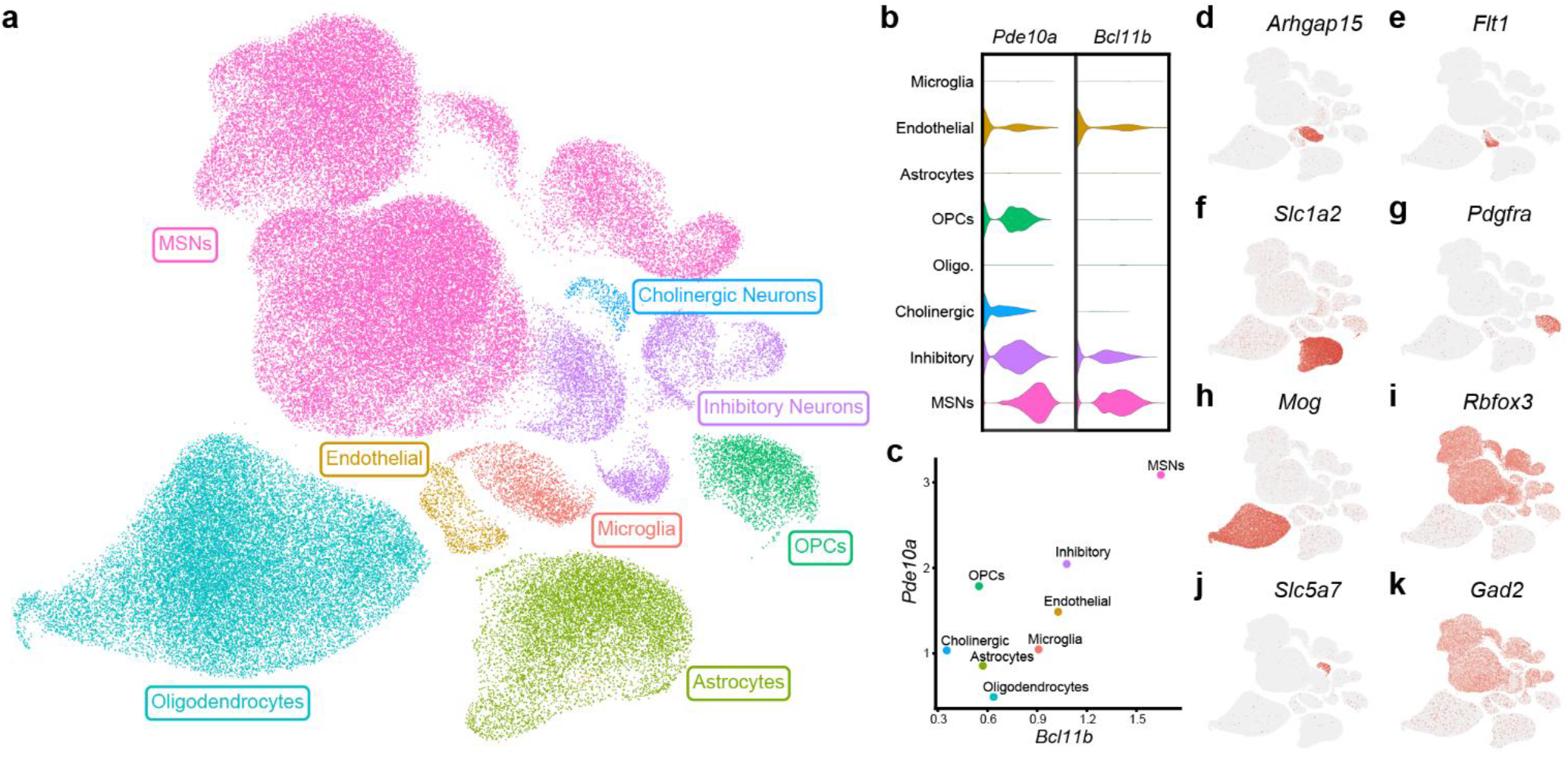
Clustering of snRNA-seq data. Single nucleus RNA sequencing was performed on nucleus accumbens samples from male, drug-naïve Brown Norway rats and data were clustered based on transcriptomic profile. (a) Uniform manifold approximation and projection (UMAP) dimension reduction plot with nuclei colored by major cell type. Normalized expression of two MSN marker genes (*Bcl11b* and *Pde10a*) is visualized by both violin plot (b) and scatter plot (c). (d-k) Feature plots display expression of markers for major glial populations and non-MSN neuronal subtypes for cluster identification.

To identify populations of MSNs in the NAc, MSNs were subset (*n*= 48,040 MSN nuclei) and clustered at low resolution using principal components derived from variably expressed genes. Five major populations of MSNs were identified (Figure 2A). Two clusters expressed markers of D2R-expressing MSNs (*Drd2*; Figure 2C) but differed in the expression patterns of additional marker genes like *Scube1* and *Stk32a* (Figure 2E and 2G). The remaining three clusters expressed markers of D1R-expressing MSNs (*Drd1*; Figure 2B) but were differentiated by expression of *Htr4, Ebf1* and *Ppm1e*, among other markers (Figure 2D, 2F and 2H). The identification of separate *Htr4*+ and *Ebf1*+ clusters of D1R-expressing MSNs is consistent with another recent snRNA-seq analysis of rat NAc ^28^. D1R-expressing *Ppm1e*+ cells have previously been labeled as *Grm8* MSNs ^28,30^ due to high expression of *Grm8* (Figure 2I). However, *Grm8* expression is also present in other D1R-expressing MSN populations (Figure 2I), whereas *Ppm1e* expression is not (Figure 2H). These data suggest that *Ppm1e* is the more robust marker for this major MSN subtype.

**Figure 2:**
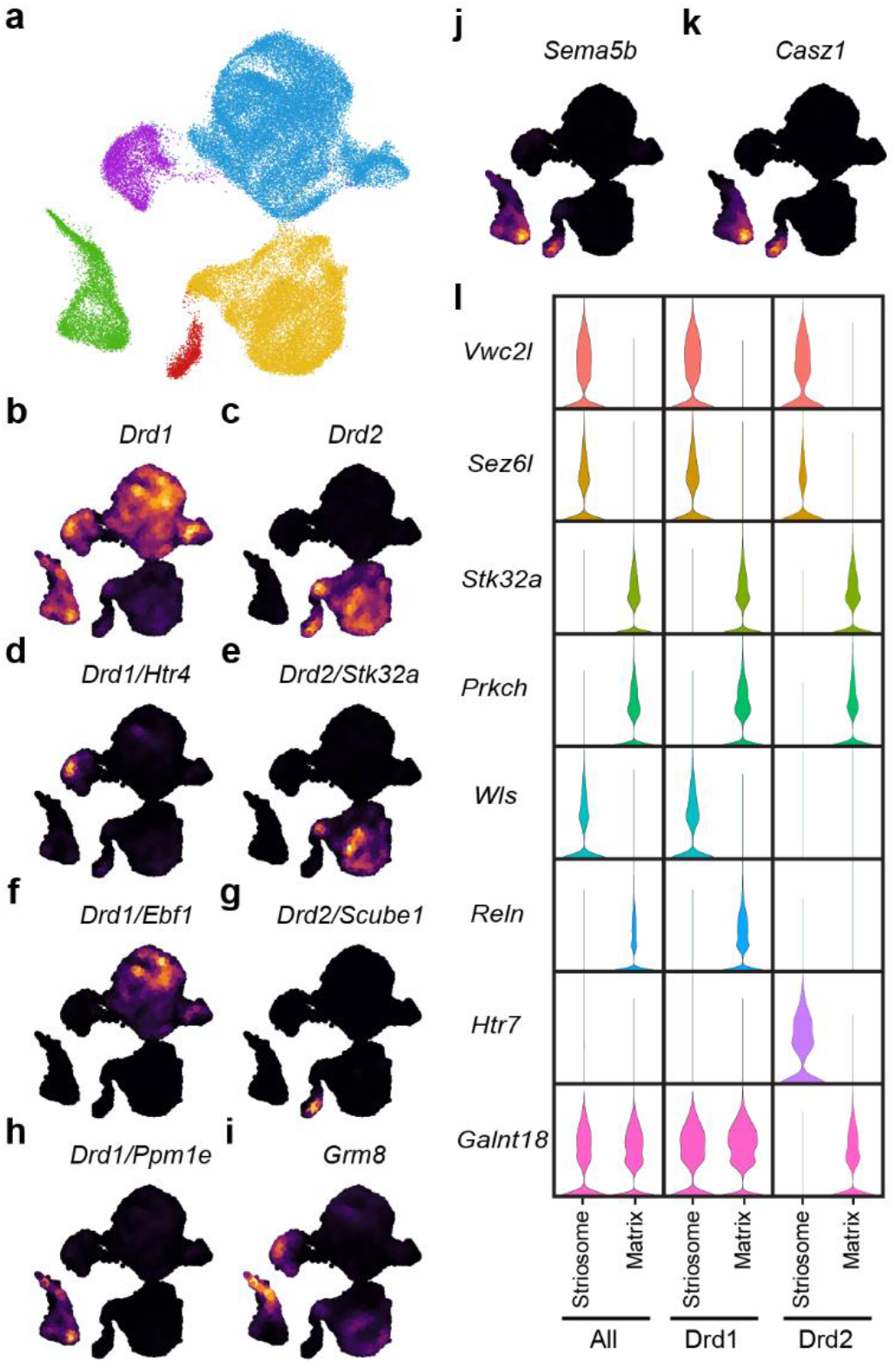
Low resolution clustering of MSNs. (a) UMAP dimension reduction plot of MSN clustered at low resolution. Nuclei are colored by major MSN population. Nebulosa plots visualizing expression on the MSN UMAP demonstrate (b) *Drd1* expression in three MSN populations and (c) *Drd2* expression in two populations, as well as (e-h) co-expression of major population markers with *Drd1* or *Drd2*. Expression of the striosomal markers (j) *Sema5b* and (k) *Casz1* is also presented. (l) Violin plots demonstrate expression differences of additional genes between all striosome or matrix MSNs, or only striosome and matrix MSNs expressing either *Drd1* or *Drd2*.

Previous single-cell analyses of mouse, macaque, and human MSNs have demonstrated that matrix and striosome neurons can be differentiated by their transcriptomic profiles ^24,26,31^. To determine whether these populations were present in our dataset, we analyzed the expression of *Sema5b* and *Casz1*, two known markers of striosome MSNs ^38,39^. Both genes were expressed in the D1R-expressing *Ppm1e*+ and D2R-expressing *Scube1*+ clusters (Figure 2J-K), indicating that these are striosome cell populations. In contrast, the *Htr4*+ and *Ebf1*+ clusters of D1R-expressing MSNs and the *Stk32a*+ cluster of D2R-expressing MSNs originate from the matrix. Comparisons of all striosome MSNs versus all matrix MSNs, D1R-expressing striosome versus D1R-expressing matrix, and D2R-expressing striosome versus D2R-expressing matrix revealed large numbers of marker genes for all groups (Figure 2L and Tables S1-3).

### Cross-species conservation of major MSN cell types

To determine whether the major MSN populations are conserved across species, we integrated our rat data with MSNs from previously published drug-naïve mouse and human NAc datasets ^25,26^. A UMAP of the integrated datasets revealed MSN clusters with contributions from all three species (Figure 3A). These clusters map to our groupings in the rat of D1R striosome (D1R *Ppm1e*+), D1R matrix (D1R *Htr4*+ and D1R *Ebf1*+), D2R striosome (D2R *Scube1*+), and D2R matrix (D2R *Stk32a*+) and the integrated data identify the analogous MSN populations in mouse and human NAc (Figure 3B-E). These findings are consistent with the previous mouse study that identified atypical, “patch-like” or striosome MSN populations expressing either *Drd1* (D1_3, D1_8) or *Drd2* (D2_2) ^26^ and demonstrate that consistent transcriptomic separation of striosome and matrix MSN cell types is possible across species.

**Figure 3:**
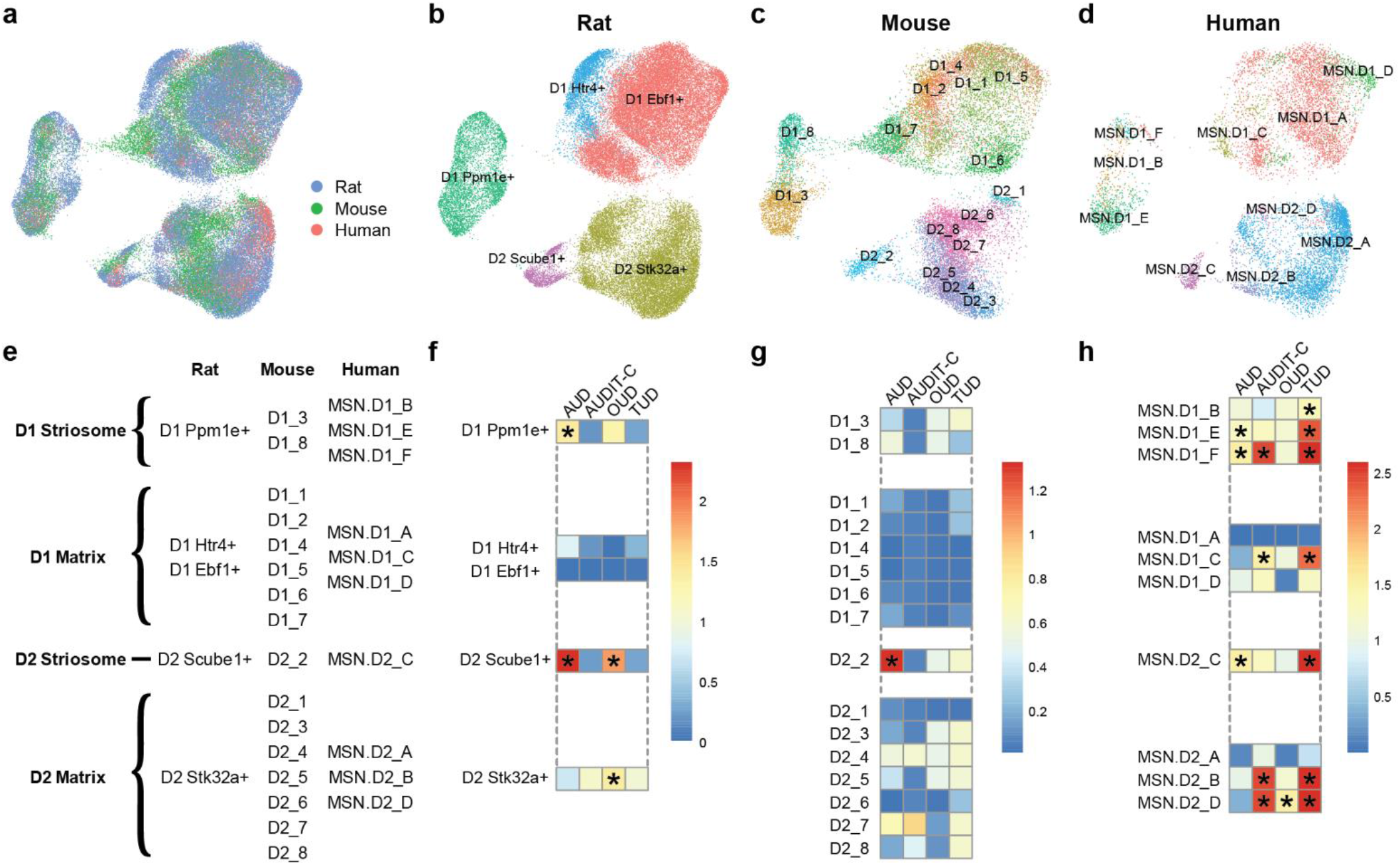
Species comparison and GWAS enrichment. MSN data from the present study were integrated with previously published MSN snRNA-seq data from mouse and human NAc. (a) A combined UMAP shows successful integration of MSNs from all three species. The MSN nuclei from each species are subsequently displayed separately and color-coded based on the cluster identifications from the current study for (b) rat and the original publications for (c) mouse and (d) human. (e) Based on the integrated data, clusters for all species are mapped to one of four major MSN populations: D1 striosome, D1 matrix, D2 striosome, and D2 matrix. Genome-wide association study summary statistics were used to assess MSN clusters in (f) rat, (g) mouse, and (h) human for enriched expression of genes associated with alcohol use disorder (AUD), alcohol consumption (AUDIT-C), opioid use disorder (OUD), and tobacco use disorder (TUD). Heatmap colors indicate the -log10(p-value) from Monte Carlo tests after correction for multiple testing with a False Discovery Rate of 0.05. Significant cell type associations are indicated by asterisks.

### Identification of MSN subtypes associated with substance use disorder phenotypes

Integrating single-cell transcriptomic data with GWAS summary statistics can implicate cell populations in phenotypes of interest and provide targets for downstream functional analyses. To identify MSN cell populations with potential relevance to SUDs, we used scDRS ^40^ to calculate disease-relevance scores for each individual cell in the rat, mouse, and human datasets using gene-level GWAS results for AUD ^37^, AUDIT-C ^37^, OUD ^36^, and TUD ^35^. The most consistent finding was a significant enrichment of genes associated with AUD in the D2R-expressing striosome clusters in all three species (Figure 3F-H). AUD-associated genes were also enriched in additional rat (D1 *Ppm1e*+) and human (MSN.D1_E, MSN.D1_F) MSN clusters, further implicating the striosome compartment in AUD (Figure 3F-H). Clusters enriched for AUD or AUDIT-C-associated genes were almost entirely non-overlapping, consistent with prior literature showing minimal genetic overlap between these two related phenotypes (Figure 3F-H) ^37,41^. D2R-expressing matrix MSNs in humans and rats, but not mice, were also significantly enriched for OUD-associated genes (Figure 3F-H). Genes associated with TUD were also enriched in 7 of the 10 human MSN clusters but no clusters in the rat or mouse MSNs (Figure 3F-H).

### Discovery of novel MSN subtypes in the rat nucleus accumbens

To identify novel MSN subtypes, principal components were derived for each of the five major populations of MSNs. Each population was then subclustered separately and the resulting datasets were merged, which resulted in 34 MSN subclusters (Figure 4A). The relationships between these new subclusters are shown in the UMAP and constellation plots in Figure 4A and 4B, respectively. Marker analyses of the 34 individual subclusters show that these populations are transcriptomically distinct (Figure 4C-G and Table S4). Although clustered at high-resolution, all subclusters can be separated and defined by the expression patterns of a relatively small number of genes (Figure 4C-G and Table S5). The subclusters produced by our high-resolution clustering include many previously undescribed subtypes of MSNs. For example, two of the novel MSN cell types, the D1R-expressing *Ppm1e*+ cells expressing *Fermt1* or *Col14a1*, are highlighted in Figure 5C and 5E, respectively. These novel subtypes each make up ∼2-3% of all MSNs in in the rat NAc. The subclustering analysis also clarified the expression patterns of prior MSN markers. We observed expression of *Chst9*, recently identified as a marker of Grm8 MSNs ^42^, in 2 of the 4 D1R *Ppm1e+* subclusters (Table S4). These data indicate that *Chst9*+ MSNs represent a distinct subset of striosomal D1R-expressing MSNs.

**Figure 4:**
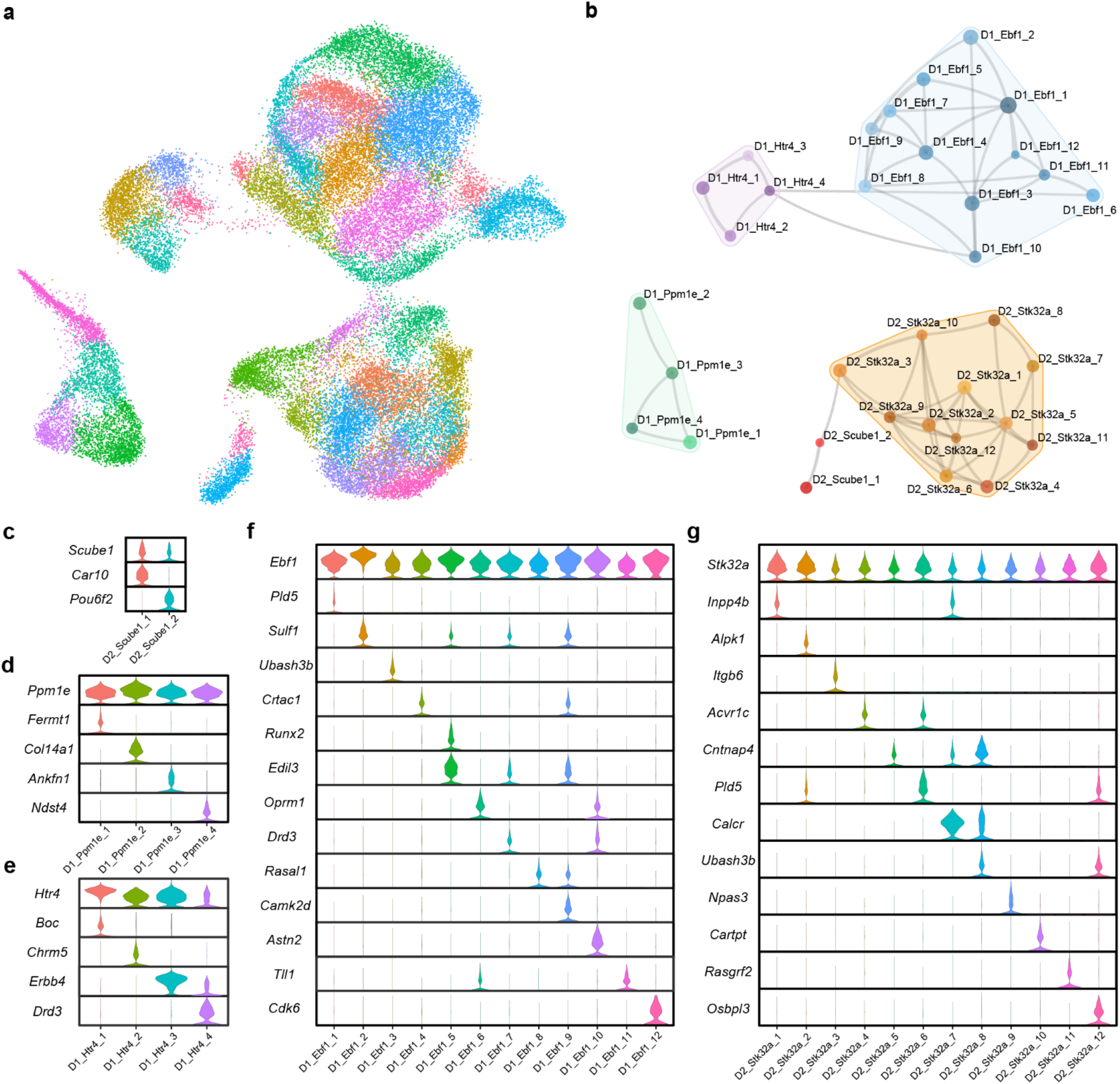
High resolution subclustering of MSNs. Each major population of MSNs was separately subclustered and then remerged. The relationships between the final 34 subclusters are displayed as both (a) a UMAP dimension reduction plot and (b) a constellation plot. Violin plots show normalized expression of marker genes for subclusters within the (c) D2 *Scube1*+, (d) D1 *Ppm1e*+, (e) D1 *Htr4*+, (f) D1 *Ebf1*+, and (g) D2 *Stk32a*+ major MSN populations.

**Figure 5:**
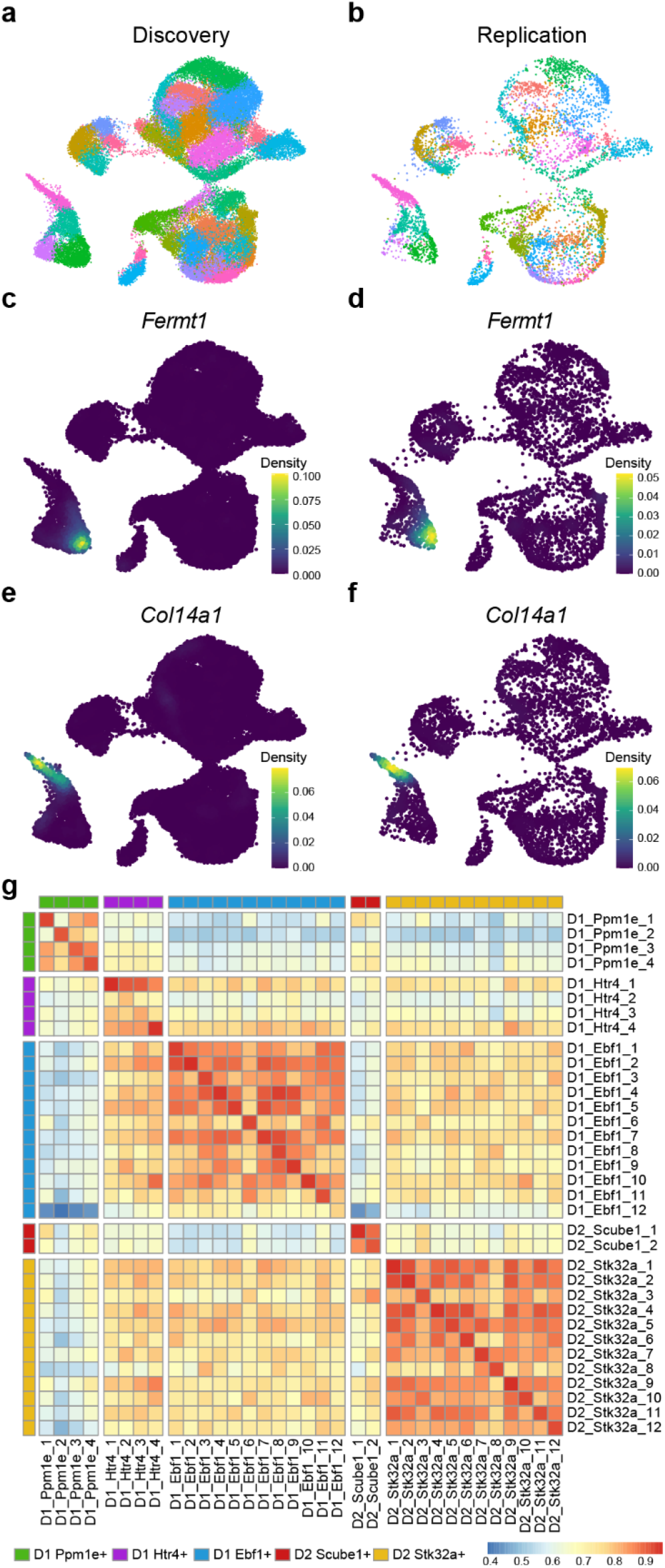
Replication of MSN subclustering in an independent snRNA-seq dataset. (a) UMAP dimension reduction plot of the current MSN dataset (n=48,040) color-coded by subcluster. (b) Rat NAc MSN nuclei (n=7,641) from previously published work ^27,28^ mapped to the discovery dataset UMAP. The replication nuclei were assigned to subclusters defined in the discovery dataset based on transcriptomic similarity and colored based on those subcluster assignments. Nebulosa plots show the expression profiles of *Fermt1* and *Col14a1* in the discovery (c,e) and replication datasets (d,f). (g) Heatmap representing Pearson correlation coefficients for gene expression between the 34 MSN subclusters in the discovery dataset and the assigned subclusters in the replication dataset.

Analysis of rat MSN subclusters with scDRS revealed several significant associations. The two D2R-expressing *Scube1*+ cell populations were significantly enriched for expression of genes associated with AUD, whereas two D2R-expressing *Stk32a*+ subclusters were enriched for genes associated with AUDIT-C score (Figure S1). No MSN cell populations were significantly associated with OUD or TUD (Figure S1).

Higher resolution clustering increases the possibility that some clusters may be driven by sample-specific variation in gene expression rather than variation between cell types, such that the clusters may not represent true cell populations. All 34 MSN subclusters contained nuclei from all samples and no sample contributed more than 25% of the nuclei in any one subcluster (Table S6), suggesting that the subclusters defined here represent distinct MSN populations in the rat NAc rather than bioinformatic artifacts.

### Profiling glutamate, GABA, and acetylcholine receptor expression in MSN subtypes

Activity of MSNs in the NAc is regulated, in part, by glutamate, GABA and acetylcholine signaling, but the expression patterns of these receptors in MSN subtypes are still not well defined. To address this knowledge gap, we analyzed the expression of genes encoding glutamate, GABA, and acetylcholine receptors in our MSN dataset (Figure S2-3). *Grik2* was the most highly expressed of all ionotropic glutamate receptor genes, with the highest expression occurring in D1R-expressing matrix MSNs, whereas *Grik3* was expressed exclusively in other MSN populations (Figure S2A). In contrast, *Grik1* had lower overall expression and was limited to a select but diverse group of MSN subclusters (Figure S2A). Expression of the metabotropic glutamate receptor gene *Grm8* also was highly variable across MSN subclusters (Figure S2B). Genes encoding GABA receptor subunits were generally less variable across the MSN cell populations, with *Gabrg3, Gabrb3*, and *Gabrb1* having notably higher expression than other family members (Figure S2C). *Chrna7* and *Chrng* were the only nicotinic acetylcholine receptors with notable expression in the dataset (Figure S3A). Several muscarinic cholinergic receptors were also expressed, with *Chrm3* showing highly variable expression and *Chrm4* only present in D1R-expressing MSNs (Figure S3B).

### Replication of novel MSN subtypes in an independent dataset

To support the validity of the subclusters identified in our analysis, we examined MSNs from an independent snRNA-seq dataset generated from drug-naïve rat NAc (*n*=7,641 MSN nuclei) ^27,28^. Mapping this replication dataset onto our discovery dataset (*n*= 48,040 MSN nuclei) yielded cells from all 34 MSN subclusters (Figure 5A-B). The novel subtypes expressing either *Fermt1* or *Col14a1* were also present in the replication sample (Figure 5C-F) with little to no background expression of either gene observed. More broadly, Pearson correlation analysis indicated strong overlap in gene expression between the matched subclusters in the discovery and replication datasets (Figure 5G). These data support the reproducibility of our findings and highlight uncommon and previously undescribed MSN cell types and the specificity of their respective primary marker genes.

## Discussion

Single-cell transcriptomic datasets are generally small due to the substantial costs of these experiments and often lack adequate numbers of cells from uncommon populations to effectively define those populations during clustering. Overcoming the sample size limitations can reveal novel cell subtypes ^43^. An analysis of ∼1.3 million cells from across the mouse cortex and hippocampal formation by the Allen Institute for Brain Science identified 364 neuronal populations, many of which were previously unidentified ^33^. Likewise, the HypoMap atlas of hypothalamus single-cell data also defined 130 subtypes from data on ∼220,000 neurons ^32^. Integration of large-scale, multimodal single-cell data, as in the BRAIN Initiative Cell Census Network’s analysis of the motor cortex, can also help to identify large numbers of neuronal populations ^34^. Our study contributes to these large-scale neural phenotyping efforts by examining the transcriptome profile of ∼79,000 NAc MSNs across rats, mice, and humans. By focusing on MSNs in the neuroanatomically restricted region of the NAc, we identified cell populations that are conserved between species and share associations with human phenotypes, suggesting that these MSN subtypes have high potential for translational studies.

We used single-cell transcriptomic data from rat NAc tissue to perform high-resolution subclustering and phenotyping of MSNs. The cell atlas described here provides marker combinations that define each of the 34 MSN populations, including novel subtypes, and distinct expression patterns for receptor genes known to be involved in MSN function. Previously, the largest study of the NAc included ∼20,000 MSNs from drug-naïve mice ^26^. That study focused primarily on clustering results with 16 MSN populations and demonstrated that these MSN subtypes had distinct spatial patterns within the NAc ^26^. The study also included a secondary, higher-resolution subclustering of MSNs. This analysis identified 57 cell populations, most of them previously undescribed in mice, and showed clearly that significant heterogeneity exists in MSNs of the NAc ^26^. Our clustering of ∼48,000 rat MSNs similarly supports the heterogenous nature of MSNs and revealed multiple novel cell subtypes in the NAc that could be defined by expression of a relatively small number of marker genes. These subclusters were not driven by individual samples and replication analysis in an independent rat NAc dataset ^27,28^ supported the validity of these subclusters. Our data also demonstrate that NAc MSNs from the striosome and matrix can be distinguished based on transcriptomic profile, consistent with data from mice and macaques ^24,26^, and that this distinction is also evident in human NAc MSN data. The replication analysis also suggests the MSN subclusters are stable across demographic variables, given the sex and strain differences between the discovery cohort (male, Brown Norway) and replication cohort (male and female, Sprague Dawley). In total, our results highlight the significant benefit of larger sample sizes for improving cell-type discovery efforts.

Although we identified 34 MSN subclusters in this study, there is no *a priori* way to know how many truly distinct cell populations are present in a given tissue. Decisions on clustering parameters are therefore driven by a combination of prior biological knowledge of cellular diversity within the tissue, the number of cells or nuclei in the dataset, and the ability to define distinct sets of cluster markers at the chosen resolution. Historically, our knowledge of uncommon neuronal subtypes in the NAc, like most brain regions, is lacking and thus cannot provide guideposts for higher-resolution analyses. Although we identified 34 MSN subclusters, there are likely additional MSN populations yet to be defined given the greater numbers of neuronal subtypes identified in significantly larger datasets of the hypothalamus, cortex, and hippocampus ^32,33^. Increases in the number of cells or nuclei from control brain tissue, potential integration of existing drug-naïve sample datasets to increase sample sizes, and collection of multi-modal data should be prioritized for all brain regions, as it will enable deep phenotyping of neural cell populations and facilitate downstream functional analyses and translational research.

Gene expression profiles vary between different cell types. For example, expression of D1R-and D2R-expressing MSN marker genes are largely confined to their respective groups in our rat NAc data, agreeing with a prior study of the mouse NAc ^26^. Notably, we did not find any ‘hybrid’ MSNs, defined by expression of both D1R and D2R, which had previously been observed in the dorsal striatum of non-human primates and humans ^24,31^. Together, these data suggest that hybrid D1R-and D2R-expressing MSNs may not be present in the ventral striatum or that this population of cells is limited to higher order primates. In addition to expression differences between cell types, individual populations of cells may also have wide transcriptomic variability associated with altered states. Activated neurons represent a classic example of this phenomenon, showing different transcriptomic patterns than inactive neurons due to the expression of immediate early genes associated with the active state. The 34 MSN subclusters defined in this study are “transcriptomically distinct”, a term that encompasses both differences in cell type and differences in cell state, and it is difficult to parse these two differences without further functional experiments. An argument can be made that two populations differing only in state would be expected to form a gradient of expression differences rather than yielding definitive markers and RNA velocity analyses may help to identify these relations between the individual subclusters. Evidence for state changes captured in snRNA-seq data could guide future work, including studies of induction of MSN plasticity.

In addition to phenotyping MSNs, analysis of our single-cell data and gene-level GWAS data highlighted cell populations enriched for the expression of genes associated with substance use-related phenotypes. The implicated MSN subtypes are potentially relevant for understanding the pathophysiology of substance use and misuse. The most significant finding is an association between the D2R-expressing *Scube1*+ MSNs and AUD. In this analysis, analogous clusters in both mouse and human MSN data were also associated with AUD, supporting the potential translational relevance of these cells in AUD. Based on the expression of known markers, these cells represent D2R-expressing MSNs originating from the striosome. Prior work supports the hypothesis that striosome MSNs are involved in reward and reinforcement learning ^44,45^ and MSNs in the striosome project to areas of the brain involved in addiction-like behaviors (e.g. ventral tegmental area) ^46^. Furthermore, manipulation of circuits targeting the striosome or matrix compartments of the striatum had differential effects on cost-benefit decision making ^47^. Mice were more likely to choose a high-reward, high-cost option over a low-reward, low-cost option after inhibition of inputs to the striosome ^47^. Despite these compelling connections, the role of the striosome in the effects of chronic alcohol exposure is unknown. Functional phenotyping of D2R-expressing striosome MSNs by other, non-sequencing methods should be prioritized to determine the role that these cells play in the relationship between the NAc and risk of AUD. Similarly, populations of D2R-expressing matrix MSNs were significantly enriched for OUD-associated genes in both humans and rats, and warrant functional follow-up studies. In contrast, enrichment for TUD-associated genes were observed in the majority of human MSN subtypes but not in rodent populations. These data could reflect notable species differences in the expression patterns of the top genes identified in the TUD GWAS and make it difficult to hypothesize whether TUD-related circuits in the NAc are conserved.

The identification of 34 MSN subclusters raises questions about the spatial location, circuit involvement, and overall function of these cell populations. The major D1R-expressing *Htr4*+ and *Ebf1*+ MSN populations observed in our analysis were previously seen in the replication dataset and shown to have significant spatial biases towards the NAc shell and core, respectively ^28^. Analyses in mice similarly showed subregion expression differences between MSN clusters ^26^. These findings suggest that the subclusters identified in the current study are also likely to have non-uniform distributions in the rat NAc. Future experiments with highly multiplexed fluorescent *in situ* hybridization (FISH) would help define the spatial profiles for each MSN subtype. Circuit tracing could also be used to define spatial distributions and study functional relevance, the ultimate goal of the type of neuronal phenotyping presented here. It is well established that MSNs in the NAc core and shell have different functions ^13-15^. Similar differences exist between neurons in the matrix and those in the striosome ^22,23^. These findings, however, are typically based on the generic classification of MSNs within those regions as either D1R-or D2R-expressing cells. Our results indicate that manipulation of all D1R-or D2R-expressing MSNs, even if restricted to only one of these NAc subregions or compartments, will inevitably affect multiple distinct MSN subtypes. It is therefore impossible to know which populations are truly relevant to any observed phenotypic effects. Cell atlases like the one presented here make it possible to define subclusters of cells and their respective marker genes, allowing more targeted genetic manipulation and functional dissection of behavior.

## Materials and Methods

### Medium Spiny Neuron Clustering

Single nucleus RNA sequencing (snRNA-seq) data from adult male, drug-naive Brown Norway rats (*n*=11) were obtained from our prior study of the NAc ^30^. Sequencing reads were aligned to the rn7.2.110 version of the rat genome (downloaded 09.13.2023) and count matrices were generated using CellRanger v7.1.0 ^48^. Seurat objects were created for all samples using *Seurat* v4.3 ^49^. Nuclei with ≤200 genes detected or ≥5% of reads from the mitochondrial genome were removed. After initial clustering, *SoupX* v1.6.2 was used to correct the count matrix for each sample for the presence of cell-free mRNA ^50^. Expression data were normalized using SCTransform while regressing out the effect of the number of unique molecular identifiers (UMIs) per nucleus, and integration was used to combine all samples. Doublets were identified with *scDblFinder* v1.12.0 ^51^. All doublets and all clusters with a majority of nuclei labeled as doublets were removed. Additional putative doublet clusters with mixed cell type markers and clusters with low average UMIs were also removed. Data for all nuclei belonging to MSN clusters were subset and used for low-resolution clustering using the first 20 principal components (PCs) and a resolution of 0.05 to identify major cell populations. To identify MSN subtypes, each major MSN population was subclustered independently with newly calculated PCs. Subclusters with poor markers were removed and the remaining 34 subclusters were merged to create the final dataset of 48,040 MSNs. A constellation plot was created using modified versions of published code ^52^ and the *scrattch*.*hicat* R package by the Allen Institute for Brain Science (https://github.com/AllenInstitute/scrattch). All other plots were generated with the *Seurat* and *ggplot2* packages.

### Cross-species Comparison

Seurat objects were generated from processed single nucleus/cell transcriptomic datasets for drug-naïve human ^25^ and mouse ^26^ NAc. Both datasets were then subset to MSNs using cluster identifications from their respective publications. Human gene symbols were converted to mouse orthologs using the *nichenetr* R package ^53^. Rat, mouse, and human MSN datasets were integrated using *Seurat* v4.3 and analogous MSN populations were identified by UMAP.

### Single Cell Disease-Relevance Score (scDRS) Analysis

We performed gene-based association testing for 4 well-powered SUD GWAS human datasets [i.e., alcohol use disorder (AUD) ^37^, alcohol consumption as measured by AUDIT-C ^37^, opioid use disorder (OUD) ^36^, and tobacco use disorder (TUD) ^35^] in FUMA v1.5.4 ^54^ using MAGMA v1.06 ^55^, which employs multiple regression models to detect multiple marker effects that account for SNP *p*-values and linkage disequilibrium (LD) between markers. All GWAS results were based on European ancestry, and the European 1000 Genomes Project phase 3 panel was used as the LD reference. A modified version of the *scDRS* program (https://github.com/martinjzhang/scDRS) was created to convert between human and rat orthologs (https://github.com/CristLab/scDRS_Rat) and used to generate disease-relevance scores for each nucleus based on gene expression profiles and Z-scores from the MAGMA analyses ^40^. Significant enrichment of expression of disease-associated genes was assessed for each major cluster in the rat, mouse, and human MSN datasets, as well as the rat MSN subclusters, by Monte Carlo analysis using the *scDRS* standard analytical pipeline ^40^. P-values were corrected for multiple testing using a False Discovery Rate of 0.05.

### Replication Analysis

Count matrices from an independent set of previously analyzed drug-naïve rat NAc snRNA-seq samples were obtained from NCBI GEO (accession numbers: GSE137763, GSE222418) ^27,28^ and analyzed with *Seurat* v4.3. Nuclei with <500 genes or >5% mitochondrial transcripts were removed as low quality, whereas nuclei with high numbers of UMI were removed as putative multiplets. After QC, data from each sample were merged into a single Seurat object, and then normalized, scaled, and clustered. MSN clusters were identified by high expression of *Bcl11b* and *Pde10a*, known markers of MSNs ^56^ and extracted from the full dataset. The replication MSN data were mapped to our discovery MSN data UMAP using the Seurat MapQuery function and predicted cluster identities were assigned to each nucleus in the replication dataset. Pearson correlation coefficients of gene expression between clusters in the discovery and replication datasets were calculated in *clustifyr* ^57^ and visualized with *pheatmap*. Nebulosa plots were generated using the *Nebulosa* R package ^58^.

## Data availability

snRNA-seq count matrices for our discovery dataset are available at the NCBI Gene Expression Omnibus (GEO) under accession number GSE263307. The cleaned Seurat objects for the full dataset and the final subclustered MSNs are also available upon request. The raw count matrices used in the replication analysis are available at GEO accession numbers GSE137763 and GSE222418. Cleaned data from mouse and human NAc can be obtained at GEO accession number GSE118020 and https://github.com/LieberInstitute/10xPilot_snRNAseq-human, respectively. Gene-level summary statistics from SUD GWAS are available from their respective groups ^35-37^.

## Code availability

All R code used for these analyses is available at https://github.com/CristLab/msn_subclustering

## Acknowledgements

The authors would like to thank Dr. Yafang Zhang, Kael Ragnini, Jennifer Ben Nathan, and Mariela Jennings for their technical contributions to this project. This work was supported in part by a State of Pennsylvania Department of Health Nonformula Tobacco Settlement Act Grant, Pharmacogenetics of Opioid Use Disorder (W.H.B.); National Institutes of Health grants R01 DA037897 (H.D.S.), R21 DA045792 (H.D.S.), R21 DA 057458 (H.D.S. & R.C.C.), R21 DA 055846 (B.C.R.), NIH/NIDA DP1DA054394 (S.S.-R.), K01 AA028292 (R.L.K.), and R01 AA030056 (H.R.K & R.L.K); Tobacco-Related Disease Research Program (TRDRP) Grant Number T32IR5226 (S.S.-R.); and Department of Veterans Affairs grant I01 BX004820 (H.R.K.). The funding sources had no role in the conceptualization of this work, analysis of the data, or preparation of the manuscript.

## Author Contributions

B.C.R., H.D.S., and R.C.C. conceptualized the project. B.C.R., W.H.B., H.R.K., S.S.-R., R.L.K., H.D.S., and R.C.C. obtained funding for the project. B.C.R., H.R.K., R.L.K., H.D.S., and R.C.C. designed the analyses.B.C.R., S.N.M., R.M., H.D.S., and R.C.C. interpreted the analyses and outcomes. B.C.R. and R.C.C. performed the bioinformatic analyses. S.T., H.R.K., S.S.-R., and R.L.K. conceptualized the human genome-wide association studies, provided genetic data and performed genetic analyses. R.C.C., B.C.R., and H.D.S. wrote the original manuscript. All authors participated in reviewing and editing the manuscript for publication.

