## Supplemental Figures 1-3 for "A single-nucleus transcriptomic atlas of medium spiny neurons in the rat nucleus accumbens"

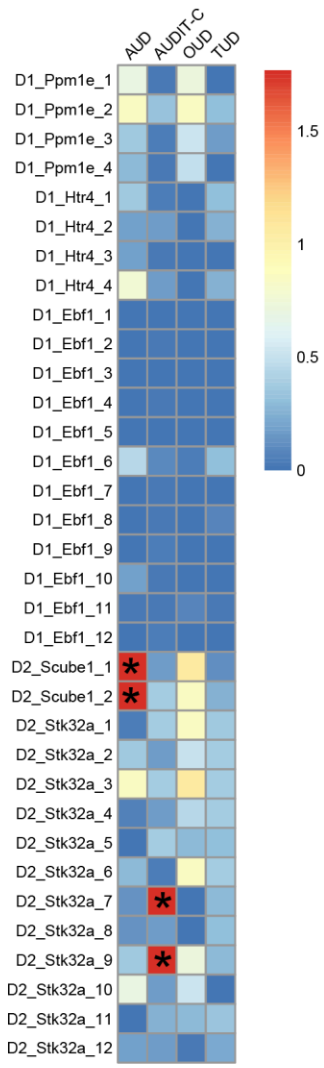


**Figure S1: MSN subcluster GWAS enrichment.** Genome-wide association study summary statistics were used to assess MSN subclusters for enriched expression of genes associated with alcohol use disorder (AUD), alcohol consumption (AUDIT-C), opioid use disorder (OUD), and tobacco use disorder (TUD). Heatmap colors indicate the -log10(p-value) from Monte Carlo tests after correction for multiple testing with a False Discovery Rate of 0.05. Significant cell type associations are indicated by asterisks.

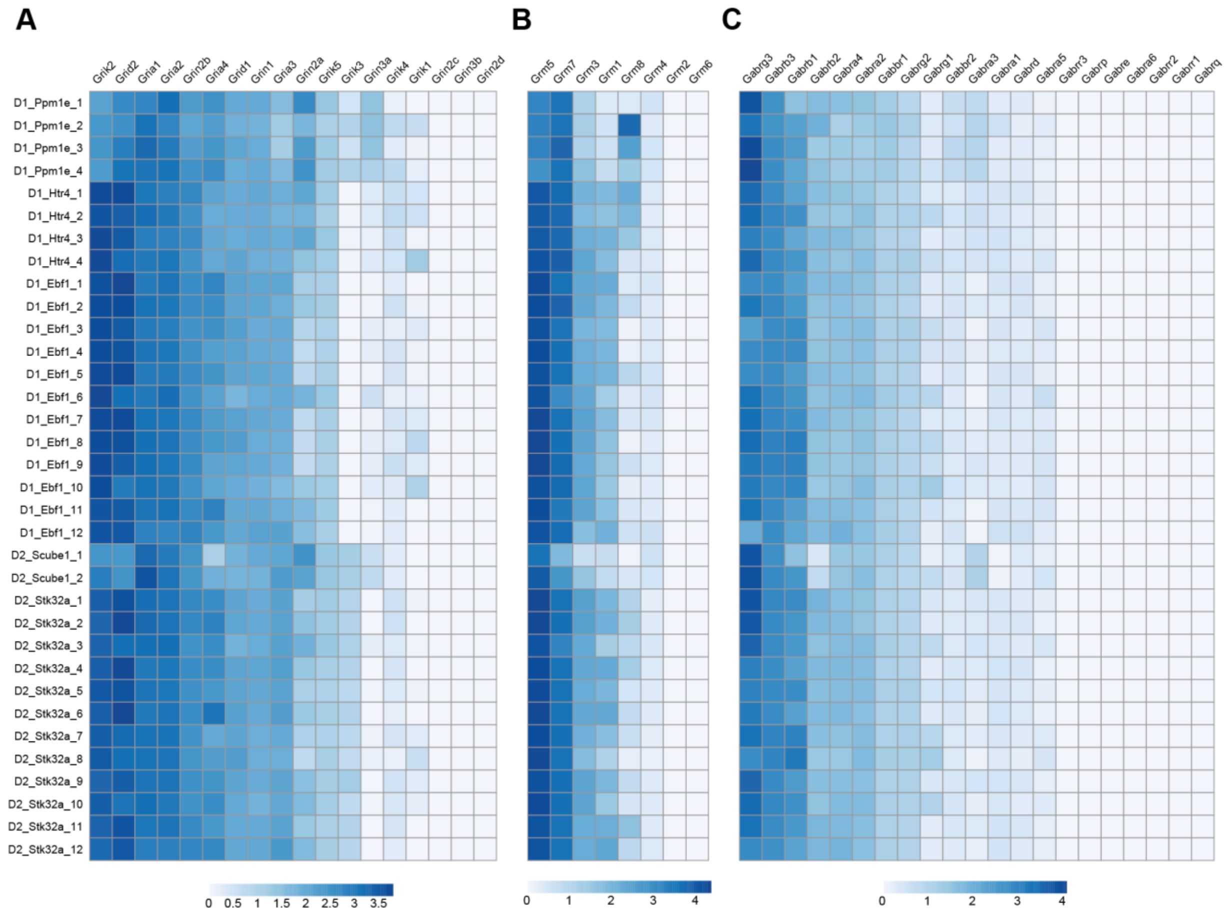


**Figure S2: Expression of glutamate and GABA receptor genes in NAc MSN subclusters.** Heatmaps indicate normalized expression of (A) ionotropic and (B) metabotrophic glutamate receptor genes and (C) GABA receptor genes in MSN subclusters from the rat NAc. Genes in each panel are sorted from highest to lowest expression average expression.

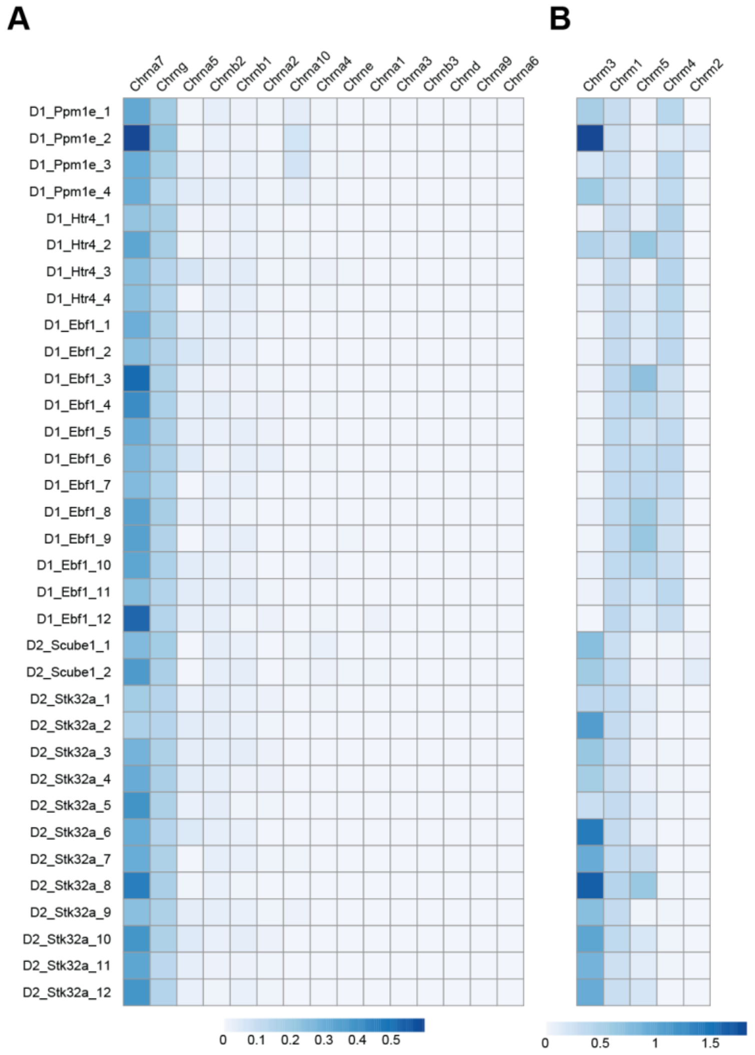


**Figure S3: Expression of acetylcholine receptor genes in NAc MSN subclusters.** Heatmaps indicate normalized expression of (A) nicotinic and (B) muscarinic acetylcholine receptor genes in MSN subclusters from the rat NAc. Genes in each panel are sorted from highest to lowest expression average expression.
